# Retrograde induction of phyB orchestrates ethylene-auxin hierarchy to regulate growth

**DOI:** 10.1101/2020.01.31.929133

**Authors:** Jishan Jiang, Yanmei Xiao, Wei Hu, Hao Chen, Liping Zeng, Haiyan Ke, Franck A. Ditengou, Upendra Devisetty, Klaus Palme, Julin Maloof, Katayoon Dehesh

## Abstract

Exquisitely regulated plastid-to-nucleus communication by retrograde signaling pathways is essential for fine-tuning of responses to the prevailing environmental conditions. The plastidial retrograde signaling metabolite methylerythritol cyclodiphosphate (MEcPP) has emerged as a stress signal transduced into a diverse ensemble of response outputs. Here we demonstrate enhanced phytochrome B protein abundance in red-light grown MEcPP-accumulating mutant (*ceh1*) plant relative to wild-type seedlings. We further establish MEcPP-mediated coordination of phytochrome B with auxin and ethylene signaling pathways, and uncover differential hypocotyl growth of red-light grown seedlings in response to these phytohormones. Genetic and pharmacological interference with ethylene and auxin pathways outline the hierarchy of responses, placing auxin epistatic to the ethylene signaling pathway. Collectively, our finding establishes the key role of a plastidial retrograde metabolite in orchestrating the transduction of a repertoire of signaling cascades, and positions plastids at the zenith of relaying information coordinating external signals and internal regulatory circuitry to secure organismal integrity.

**Two sentence summary:** The plastidial retrograde metabolite, MEcPP, orchestrates coordination of light and hormonal signaling cascade through induction of phytochrome B abundance and modulation of auxin and ethylene levels for optimal adaptive responses to light environment.

## Introduction

Dynamic alignment of internal and external cues through activation of corresponding signal transduction pathways is a defining characteristic of organisms essential for fitness and the balancing act of metabolic investment in growth versus adaptive responses. The integrity of these responses is achieved through finely controlled communication circuitry, notably retrograde (organelle-to-nucleus) signaling cascades. Despite the central role of retrograde signaling in regulation and coordination of numerous adaptive processes, the nature and the operational mode of action of retrograde signals have remained poorly understood.

Through a forward genetic screen, we identified a bifunctional plastid-produced metabolite methylerythritol cyclodiphosphate (MEcPP) that serves as a precursor of isoprenoids produced by the plastidial methylerythritol phosphate (MEP) pathway and functions as a stress-specific retrograde signaling metabolite (Xiao et al., 2012). We further demonstrated that stress-induced MEcPP accumulation leads to growth retardation and induction of selected nuclear encoded stress response genes (Xiao et al., 2012; Walley et al., 2015; Lemos et al., 2016; Wang et al., 2017a). We specifically established that regulation of growth is in part via MEcPP-mediated modulation of levels and distribution patterns of auxin (IAA) through dual transcriptional and post-translational regulatory inputs (Jiang et al., 2018).

Auxin functions as a key hormone regulating a repertoire of plant development processes including hypocotyl growth (J. Jensen et al., 1998; De Grauwe et al., 2005). The auxin biosynthesis pathway that converts tryptophan (Trp) to IAA in plants is established to be through conversion of Trp to indole-3-pyruvate (IPA) by the TAA family of amino transferases and subsequent production of IAA from IPA by the YUC family, a family of flavin monooxygenases (Zhao, 2012). Subsequently, establishment of auxin gradient is achieved by transporters such as the auxin-efflux carrier PIN-FORMED1 (PIN1) (Galweiler et al., 1998; Geldner et al., 2001). Interestingly, IAA biosynthesis, transport, and signaling during light-mediated hypocotyl growth is in turn regulated by ethylene (Liang et al., 2012), and conversely ethylene is regulated by auxin (Vandenbussche et al., 2003; Ruzicka et al., 2007; Stepanova et al., 2007; Swarup et al., 2007; Negi et al., 2010). As such, auxin-ethylene crosstalk inserts an additional layer of complexity to the already intricate and multifaceted growth regulatory mechanisms.

Ethylene in plants is derived from conversion of S-adenosyl-L-methionine (AdoMet) to 1-aminocyclopropane-1-carboxylate (ACC) by ACC synthase (ACS) (Yang and Hoffman, 1984), followed by conversion of ACC to ethylene catalyzed by ACC oxidase (L.-C. Wang et al., 2002). Ethylene stimulates hypocotyl growth in the light but inhibits it in the dark (Smalle et al., 1997; J. Jensen et al., 1998; Vandenbussche et al., 2012).

Light signaling is a common environmental stimulus controlling developmental processes through hormonal modulation, such as regulation of auxin biosynthesis and signaling genes by phytochrome B (phyB) (Morelli and Ruberti, 2002; Tanaka et al., 2002b; Tian et al., 2002; Nozue et al., 2011; Hornitschek et al., 2012a; de Wit et al., 2014; Leivar and Monte, 2014). PhyB is the main photoreceptor mediating photomorphogenesis by red light; phyB is activated by red light and imported into the nucleus where it forms phyB-containing nuclear bodies (phyB-NBs) (Nagy and Schafer, 2002; Quail, 2002). Formation of phyB-NBs depends on binding to and sequestration of the basic helix-loop-helix (bHLH) transcription factors, Phytochrome Interacting Factor 1 (PIF1), PIF3, PIF4, PIF5, and PIF7 (Rausenberger et al., 2010; Leivar and Quail, 2011). The prominent role of phyB in auxin regulation is best displayed by simulation of shade avoidance responses (SAR) through exogenous application of auxin or via genetic manipulation of auxin (Tanaka et al., 2002a; Hornitschek et al., 2012b). In addition, PIFs, specifically *PIF4, PIF5, PIF7* play a major role in regulating auxin by targeting promoter elements of multiple auxin biosynthetic and signal transduction genes (Franklin et al., 2011; Nozue et al., 2011; Leivar et al., 2012; Sellaro et al., 2012; Leivar and Monte, 2014). Moreover, the ethylene-promoted hypocotyl elongation in light is regulated by the PIF3-dependent growth-promoting pathway activated transcriptionally by EIN3; while under dark condition, ethylene inhibits growth by destabilizing the ethylene response factor 1 (ERF1) (Zhong et al., 2012).

Here we have identified MEcPP as the retrograde signaling metabolite that coordinates internal and external cues, and we further delineated light and hormonal signaling cascades that elicit adaptive responses to ultimately drive growth-regulating processes tailored to the prevailing environment.

## Results

### Elevated phyB abundance suppresses hypocotyl growth in ceh1

Given the stunted hypocotyl phenotype of high MEcPP-accumulating mutant plant (*ceh1*), we explored the nature of the photoreceptor involved by examining hypocotyl length of seedlings grown in the dark and under various monochromatic light conditions. The analyses showed comparable hypocotyl lengths of the dark-grown *ceh1* and the control seedlings (WT) (Fig. 1**A**). However, under continuous red light (Rc), *ceh1* seedlings displayed notably shorter hypocotyls than those of WT plants (Fig. 1**A**). This data led us to question the role of phyB, the prominent red light photoreceptor, in regulating *ceh1* hypocotyl growth. To answer this question, we generated a *ceh1/phyB-9* double mutant line, and subsequently compared their hypocotyl length with those of WT, *ceh1* and *phyB-9* seedlings grown under continuous dark and Rc conditions (Fig. 1**A** & **B**). The data clearly demonstrate phyB-dependent suppression of hypocotyl growth in *ceh1* under Rc, as evidenced by the recovery of *ceh1* retarded hypocotyl growth in the *ceh1/phyB-9* to comparable lengths as those of *phyB-9* seedlings.

**Fig. 1.**
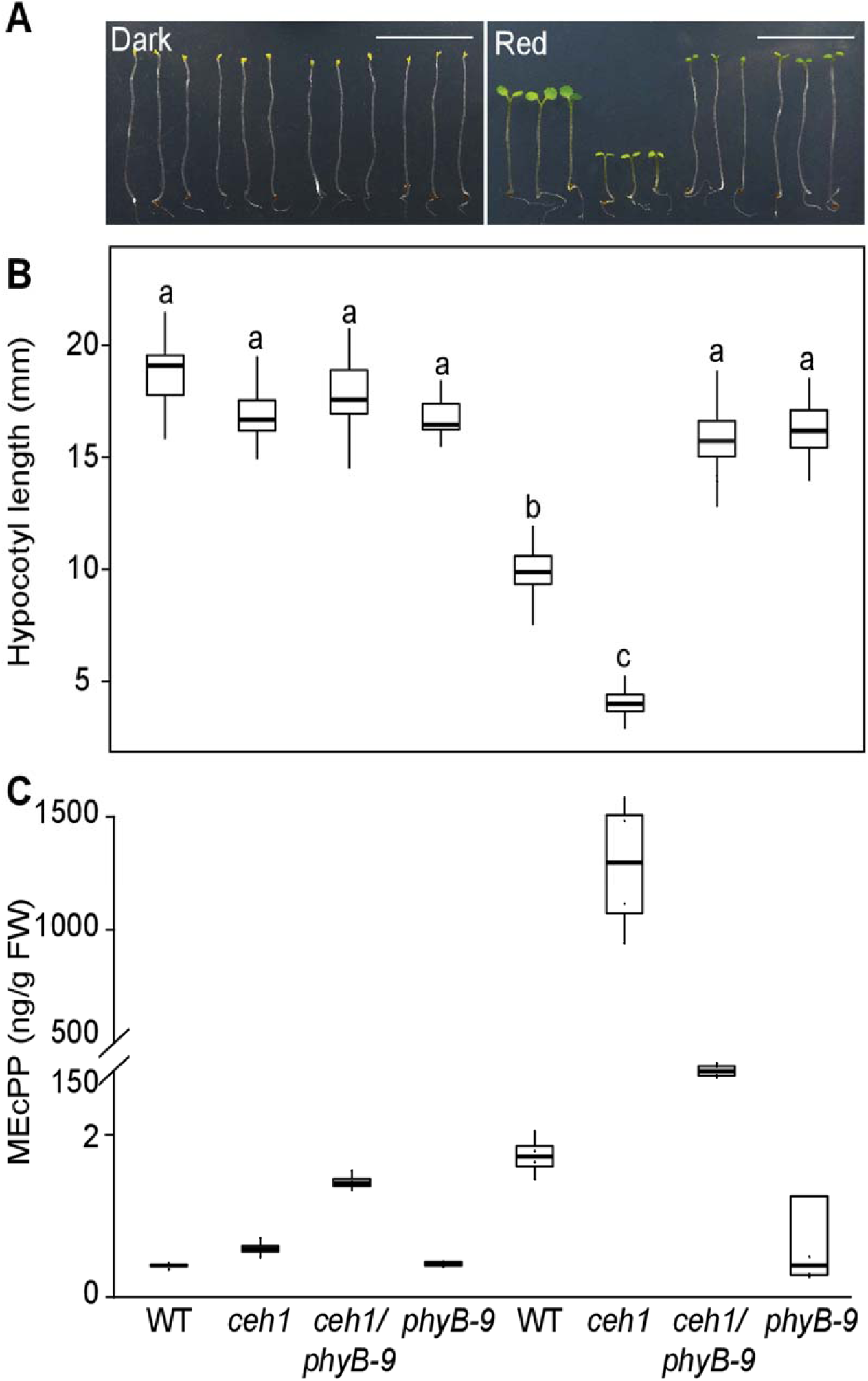
*Ceh1* hypocotyl growth in red light is phyB-dependent. (**A**) Representative images of 7-day-old WT, *ceh1, ceh1/phyB-9* and *phyB-9* seedlings grown in the dark and continuous red light (Rc: μEm^−2^sec^−1^). (**B**) Quantification of hypocotyl lengths from aforementioned genotypes shown in Fig. **1 A**. (**C**) MEcPP levels of samples from Fig.**1A**. Statistical analyses were performed using Tukey’s HSD method, different letters indicate significant difference (*P* < 0.05). Scale bars: 1cm.

Hypocotyl growth of the aforementioned four genotypes was also examined under continuous blue (Bc) and far red (FRc) light conditions. The reduced hypocotyl growth of *ceh1* mutant grown under Bc, albeit not as strongly as those grown under Rc, further implicate blue light receptor, cytochrome, (Yu et al., 2010) in regulating growth of these seedlings (Fig. S1**A** & **B**). Additionally, *ceh1* and *ceh1/phyB-9* seedlings grown under Bc exhibited equally shortened hypocotyls, and under FRc light the growth was almost similarly retarded in all genotypes (Fig. S1**A** & **B**). Collectively, these results support involvement of cryptochrome as well and phyB in *ceh1* hypocotyl growth, albeit at different degrees. However, the more drastic effect of phyB in regulating hypocotyl growth of Rc grown high MEcPP accumulating seedlings in conjunction with the supporting evidence from the earlier data using white light grown *ceh1* seedlings (Jiang et al., 2019), led us to primarily focus on the role of phyB.

Next, we measured MEcPP levels in the four genotypes grown in the dark and various monochromatic wave lengths to examine a potential correlation between growth phenotypes and altered levels of the retrograde signaling metabolite (Fig. 1**C** & S1**C**). The analyses showed almost undetectable MEcPP levels in all the dark grown genotypes, and low levels of the metabolite in the Rc-grown WT and *phyB-9* seedlings. In contrast, *ceh1* seedlings grown in Rc accumulated high MEcPP levels, a phenotype that was partially (∼10-fold) suppressed in *ceh1/phyB-9* seedlings. This reduction was not unexpected since phyB controlled PIF regulates the expression of *DXS*, the first MEP-pathway gene encoding the flux determinant enzyme (Chenge-Espinosa et al., 2018). It is noteworthy that despite this significant reduction, the MEcPP content of *ceh1/phyB-9* seedlings remained ∼100-fold above those of WT or *phyB-9* plants grown simultaneously and under the same conditions. This reduction of MEcPP in *ceh1*/*phyB-9* also occurred in seedlings grown in Bc (Fig. S1**C**), likely as the result of the direct interaction between PIFs and blue light receptor cryptochromes (Pedmale et al., 2016). However, in spite of reduced MEcPP levels in Rc or Bc grown *ceh1/phyB-9* seedlings, the hypocotyl growth recovery is exclusive to mutant seedlings grown in Rc (Fig. 1**A**-**C** & S1**A**-**C**). Moreover, hypocotyls of all genotypes, regardless of their MEcPP levels, remain stunted in FRc, known to inactivate phyB. Collectively, the result further verifies the functional of phyB in altering the observed growth phenotype of *ceh1* mutant seedlings.

Next, we questioned whether phyB transcript and/or protein levels are altered in *ceh1* mutants grown in Rc. While expression data analyses display similar *PHYB* transcript levels in *ceh1* and the WT seedlings (Fig. S1**D**), immunoblot analyses show notable increase in phyB protein abundance in the *ceh1* mutant compared to the WT seedlings (Fig. S1**E** & S1**F**), verifying the earlier report using white light grown seedlings (Jiang et al., 2019).

To examine the correlation between accumulation of MEcPP and alteration of growth and phyB protein abundance in response to red light treatment, we further examined the hypocotyl length of wild type (Col-0 ecotype), P (Col-0 transformed with *HPL:LUC* construct), *ceh1* and complemented *ceh1* (CP) seedlings (Fig. 2**A**-**C**). This data clearly show recovery of *ceh1* stunted hypocotyl in CP at comparable length to those of Col-0 and P seedlings (Fig. 2**A**-**B**). In addition, immunoblot analyses show similar phyB levels in Col-0, P and CP compared to the increased abundance in *ceh1* (Fig. 2**C**). Lastly, similarity of hypocotyl length and phyB abundance phenotypes between P and Col-0 seedlings clearly support suitability and wild type functionality of P genotype for additional analyses.

**Fig. 2.**
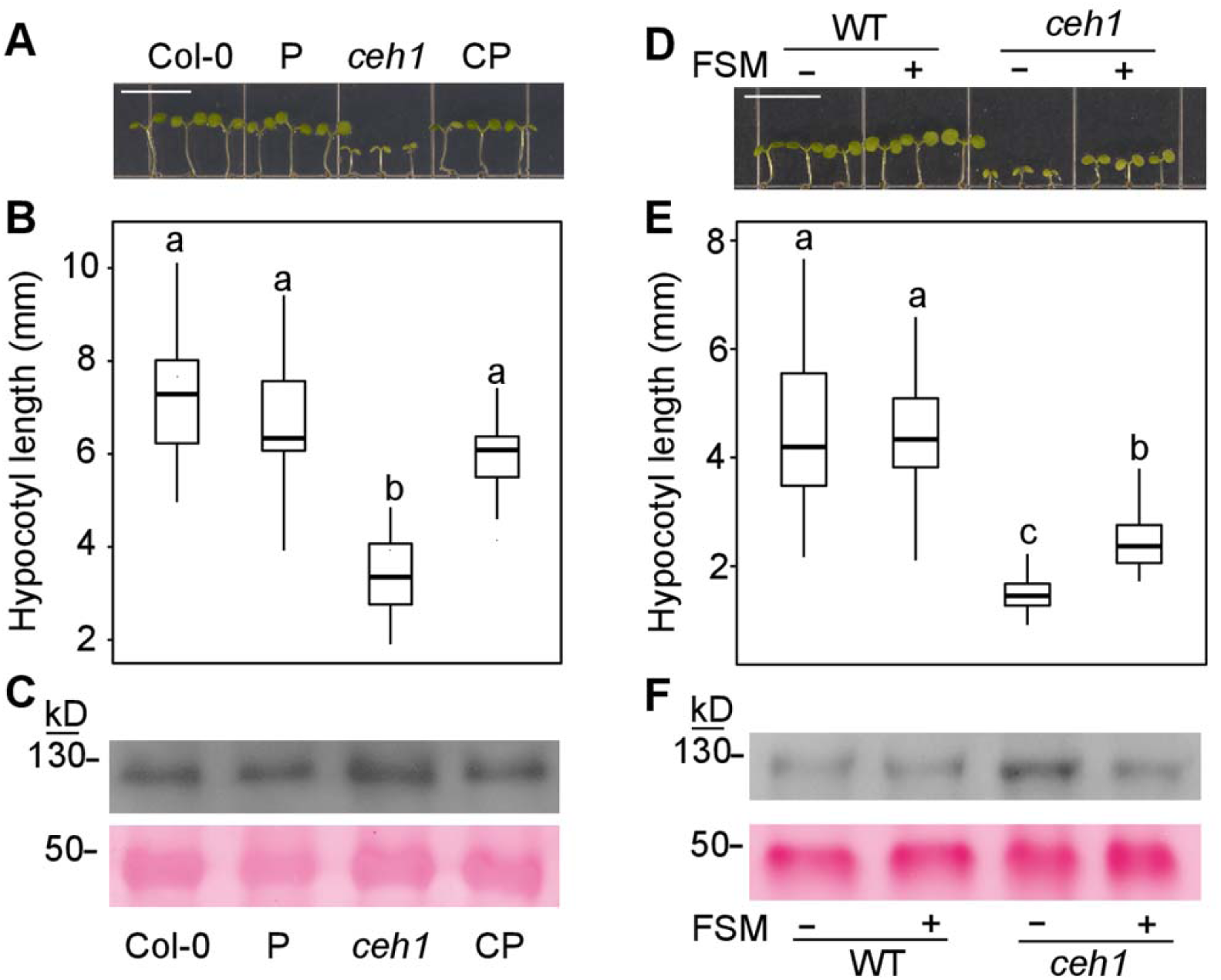
MEcPP induction of phyB results in stunted *ceh1* hypocotyl growth. (**A**) Representative images of 7-day-old Col-0, P, *ceh1* and complementation line (CP) seedlings grown in Rc (μEm^−2^sec^−1^). Scale bars: 1 cm. (**B**) & (**E**) Quantification of hypocotyl length of seedlings from panel (**A**) and (**D**), respectively. Data are presented with 45 seedlings. Statistical analyses were carried out using Tukey’s HSD method, different letters indicate significant difference (*P*< 0.05). (**C**) & (**F**) phyB protein abundance from panel (**A**) and (**D**), respectively. (**D**) Representative images of 7-day-old WT and *ceh1* seedlings grown in Rc (μEm^−2^sec^−1^) in the absence (-) and presence (+) of fosmidomycin (20 μM).

To further examine the potential role of MEcPP in *ceh1* mutant in reducing growth and altering phyB levels, we employed a pharmacological approach using fosmidomycin (FSM), a MEP-pathway inhibitor (Fig. 2**D**-**F**). This inhibitor interferes with and highly reduces the flux through the pathway and abolishes MEcPP-mediated actions such as formation of otherwise stress-induced subcellular structures known as ER bodies or enhancing the reduced auxin levels in *ceh1* mutant plants (Gonzalez-Cabanelas et al., 2015; Wang et al., 2017b; Jiang et al., 2018). We examined hypocotyl growth of red-light grown 7-day-old seedlings that were treated with FSM for 3 days. This data shows enhanced hypocotyl growth of FSM treated *ceh1* compared to non-treated seedlings (Fig. 2**D**-**E**). It is of note that the length of FSM treated *ceh1* hypocotyls have not recovered to that of the WT seedlings, suggesting inefficiency of FSM treatment and/or presence of other regulatory factors. Regardless, the immunoblot shows reduced phyB abundance in FSM treated *ceh1* compared to untreated seedling (Fig. 2**F**), supporting the notion of MEcPP-mediated increase of phyB abundance.

In addition to MEcPP, the *ceh1* mutants accumulate substantial amounts of the defense hormone, salicylic acid (SA) (Xiao et al., 2012; Bjornson et al., 2017). The reported involvement of phyB in SA accumulation and signaling (Chai et al., 2015; Nozue et al., 2018) prompted us to examine the potential role of this defense hormone in regulation of *ceh1* hypocotyl growth. For these experiments we employed previously generated SA deficient double mutant line, *ceh1*/*eds16* (Xiao et al., 2012). All four genotypes (WT, *ceh1, ceh1/eds16* and *eds16*) displayed similar hypocotyl length when grown in the dark, while in Rc both *ceh1* and *ceh1/eds16* seedlings display equally shortened hypocotyl as compared to their respective backgrounds (Fig. S1**G**). The result illustrates SA-independent regulation of *ceh1* hypocotyl growth in Rc.

Given the well-established role of PIFs in transduction of phyB signals, we examined *PIF*s expression levels and found significantly reduced *PIF4* and *5* transcripts in the Rc-grown *ceh1* compared to the WT seedlings (Fig. S2). This data led us to genetically investigate the potential role of PIFs in regulating hypocotyl length of Rc grown *ceh1* seedlings. For these experiments we quantified hypocotyl growth of *pifq (pif1, 3, 4* and *5)* alone and in lines introgressed into the *ceh1* mutant background. The results show similarly dwarf hypocotyls in *ceh1*/*pifq* and *pifq* backgrounds, which are slightly but significantly shorter than that of the *ceh1* (Fig. 3**A**). Furthermore, equally reduced hypocotyl growth in *ceh1*/*pifq* and *pifq*, suggest that PIFs are the predominant growth regulators in *ceh1* under the experimental conditions employed. The role of PIFs in determining hypocotyl growth was further tested by examining *ceh1* overexpressing *PIF 4* and *5* seedlings grown in Rc (Fig. 3**B**). The data show the expected enhanced hypocotyl growth of *PIF* over-expressers compared to the WT seedlings and the recovery of *ceh1* retarded growth in *ceh1*/*PIF 4* and *5* overexpression lines.

**Fig. 3.**
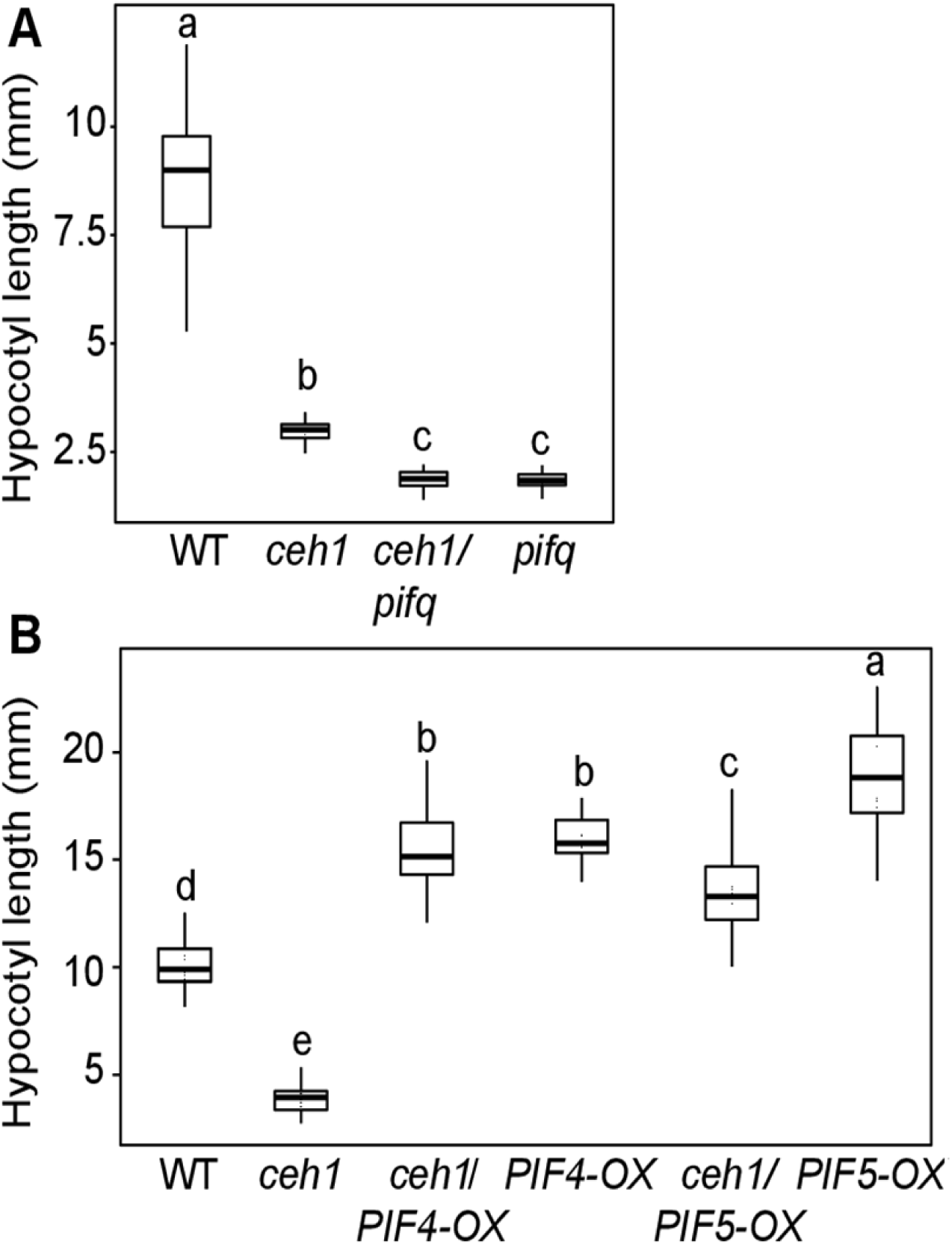
Overexpression of *PIF4* and *PIF5* recover stunted hypocotyl growth of *ceh1*. (**A**) Quantification of hypocotyl lengths of 7-day-old WT, *ceh1, ceh1/pifq*, and *pifq* grown in Rc (15 μEm^−2^sec^−1^). (**B**) Quantification of hypocotyl lengths from 7-day-old WT, *ceh1, ceh1/PIF4-OX, PIF4-OX, ceh1/PIF5-OX* and *PIF5-OX*grown in Rc (15 μEm^−2^sec^−1^). Data are presented with n≥20 for the *pif* mutant backgrounds and n≥30 for the experiments containing *PIF-OX* seedlings. Statistical analyses were carried out using Tukey’s HSD method, different letters indicate significant difference *(P*< 0.05).

Collectively this data illustrates growth regulatory function of PIFs, and identifies MEcPP-mediated transcriptional regulation of *PIF4* and *5* as an integral regulatory circuit controlling *ceh1* hypocotyl growth.

### Reduced expression of auxin biosynthesis and response genes in ceh1

To identify the downstream components of the MEcPP-mediated phyB signaling cascade, we performed RNAseq profiling of WT and *ceh1* seedlings grown in the dark and in Rc (15 μE m^−2^sec^−1^). A multi-dimensional scaling (MDS) plot revealed significant overlap between expression profiles of WT and *ceh1* seedlings grown in the dark, in contrast to their distinct expression profiles when grown in Rc (Fig. S3). GO-term analyses identified over-representations of auxin biosynthesis and response genes amongst the significantly (≥2-fold) altered transcripts (Fig S4). Confirmation of the data using qPCR identified auxin biosynthesis (*YUC3* and *8*) and response genes (*IAA6* and *19*) as the most significantly differentially expressed genes under Rc condition (Fig. 4**A-B**). We further quantified the IAA content in plants and found similar auxin levels in dark-grown genotypes in contrast to significantly reduced levels (50%) in Rc-grown *ceh1* versus WT plants (Fig. 4**C**). We validated this finding by testing Rc grown WT and *ceh1* lines expressing the auxin signaling reporter *DR5*-GFP (Jiang et al., 2018). The reduced GFP signal in *ceh1* was on par with lower IAA levels in the mutant compared to the WT seedling (Fig. 4**D**).

**Fig. 4.**
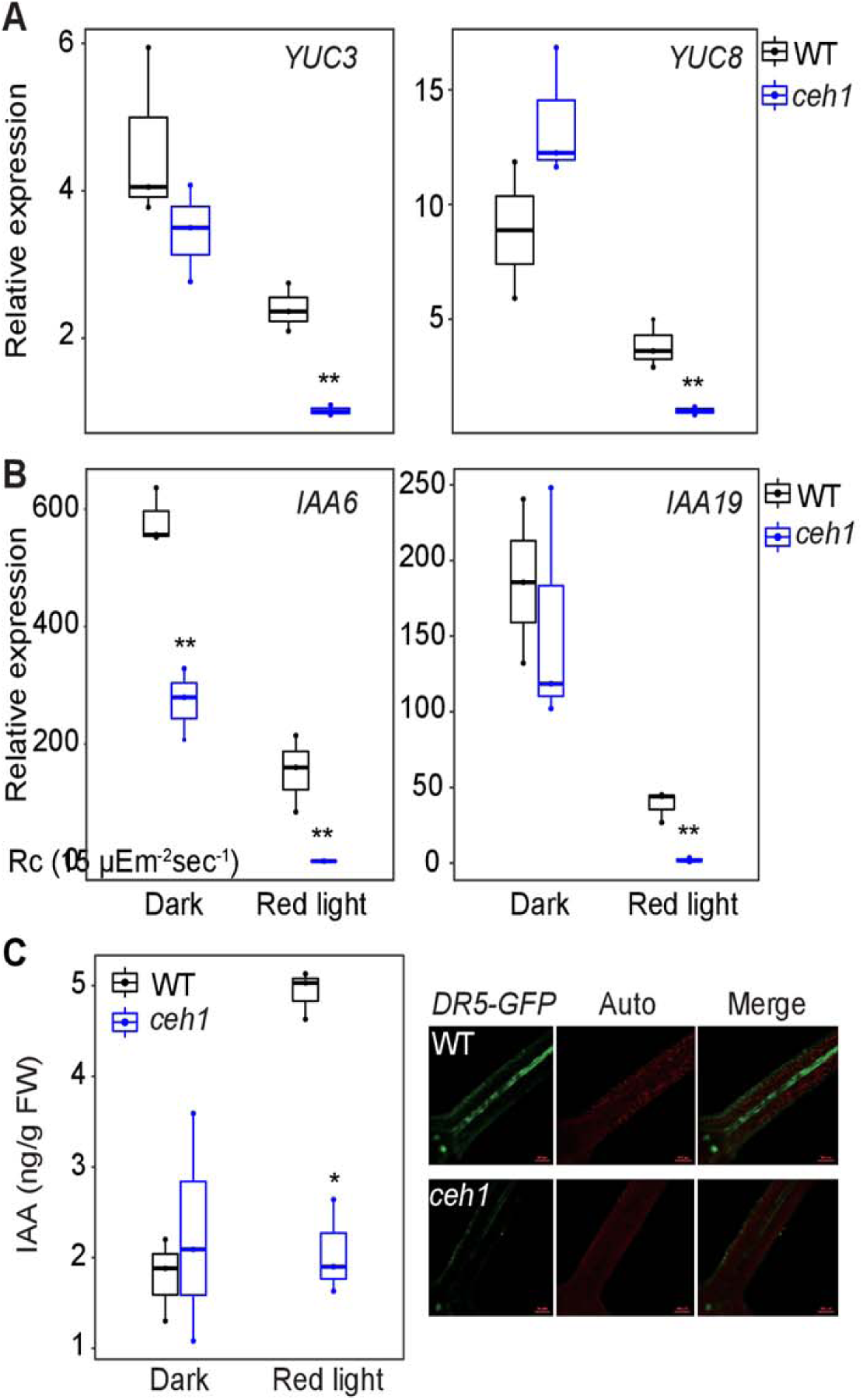
Auxin is reduced in *ceh1*. Expression levels of *YUC3, 8* **(A)** and *IAA6, 19* **(B)** in WT and *ceh1* seedlings. RNAs were extracted from 7-day-old WT and *ceh1* seedlings grown in the dark and Rc (15 μEm^−2^sec^−1^). Transcript levels of target genes were normalized to the levels of At4g26410 (M3E9). Data are presented with three biological replicates and three technical replicates. Statistical analyses were determined by a two-tailed Student’s *t* tests with a significance of *P*< 0.05 *\*, P*< 0.01 **. (**C**) IAA levels in 7-day-old WT and *ceh1* seedlings grown in the dark and Rc (15 μEm^−2^sec^−1^). Data are presented with three biological replicates. Statistical analyses were carried out by a two-tailed Student’s *t* tests with a significance of *P* < 0.05. (**D**) Representative images of *DR5-GFP* signal intensity in 7-day hypocotyls of Rc (15 μEm^−2^sec^−1^) grown WT and *ceh1* seedlings. *DR5-GFP* (green), chloroplast fluorescence (red) and merged images.

Next, we examined possible modulation of other phytohormones such as abscisic acid (ABA) and jasmonic acid (JA) in response to high MEcPP levels in *ceh1* seedlings (Fig. S**5**). Similar ABA and JA levels found in WT and *ceh1* plants grown in the dark and in Rc, strongly support the specificity of MEcPP-mediated regulation of auxin.

### Enhanced tolerance of ceh1 to auxin and auxinole

Reduced IAA levels in *ceh1* led us to examine whether external application of this hormone could rescue *ceh1* retarded hypocotyl growth. The analyses show longer hypocotyl in *ceh1* seedlings treated with IAA at growth inhibiting concentrations (10 and 100 μM) in WT seedlings (Fig. 5**A**-**B**). Interestingly, *ceh1* and WT hypocotyls displayed similar lengths when treated with the highest IAA concentration used here (100 μM), albeit through two opposing responses, namely growth suppression in WT and induction in *ceh1*.

**Fig. 5.**
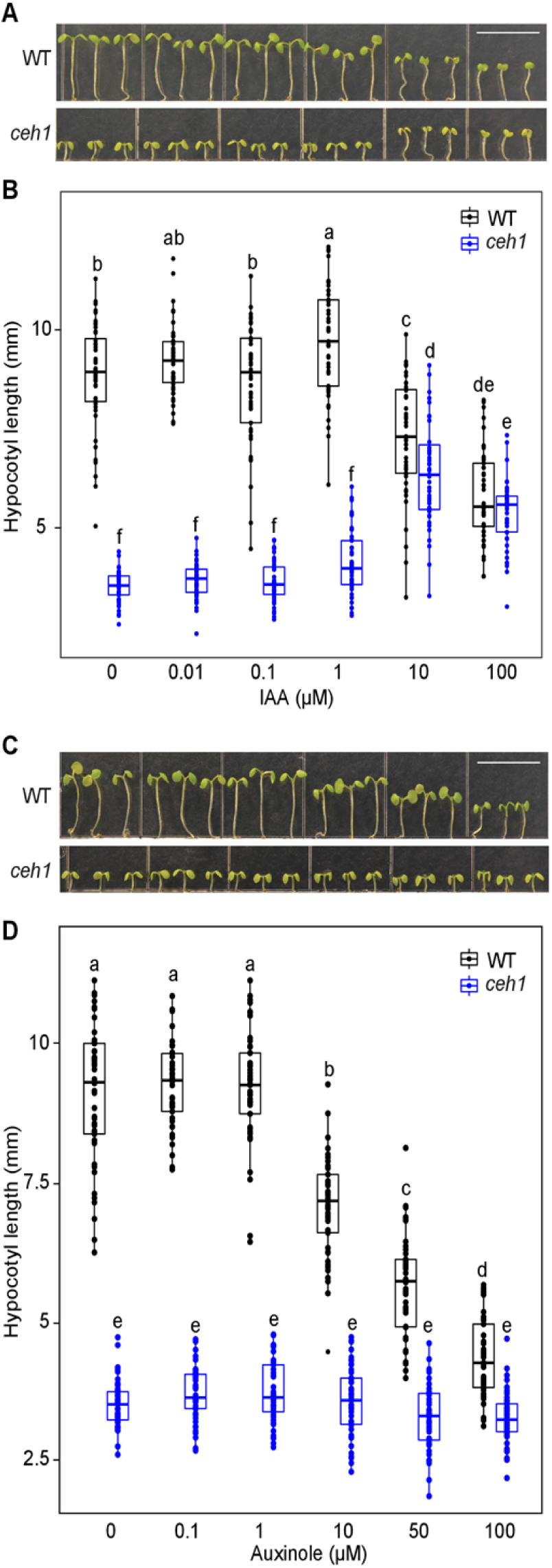
Enhanced tolerance of ***ceh1*** to auxin and auxinole. (**A**) & (**C**) Representative images of 7-day-old WT and *ceh1* seedlings in the absence (0) and presence of IAA and auxinole grown under Rc (15 μEm^−2^sec^−1^), respectively. (**B**) & (**D**) Quantification of hypocotyl lengths of seedlings from panel (**A**) & (**C**), respectively, Data are presented with 45 seedlings. Statistical analyses were carried out using Tukey’s HSD method, different letters indicate significant difference (*P* < 0.05). Scale bars: 1cm.

This finding led to the hypothesis that the enhanced tolerance of *ceh1* to auxin treatment is not solely the result of reduced auxin levels in the mutant, but also a consequence of modified auxin signaling in the mutant. To address this possibility, we treated WT and *ceh1* seedlings with auxinole, an auxin signaling inhibitor that functions as an auxin antagonist for TIR1/AFB receptors (Hayashi et al., 2008; Hayashi et al., 2012). The analyses show clear dose-dependent suppression of hypocotyl growth of WT seedlings in response to auxinole treatment, in contrast to unresponsiveness of *ceh1* seedlings at all concentrations examined (Fig. 5**C**-**D**). Collectively, the data display enhanced tolerance of *ceh1* to otherwise inhibitory concentrations of auxin and auxinole, likely stemming from reduced auxin levels and compromised signaling in the mutant line.

### Altered auxin transport in ceh1

We have previously established that MEcPP-mediated modulation of levels and distribution patterns of auxin (IAA) is via dual transcriptional and post-translational regulatory inputs (Jiang et al., 2018). We specifically demonstrated reduced transcript and protein levels of auxin efflux transporter PIN-FORMED 1 (PIN1) in *ceh1* seedlings grown in white light. Here, we extended these analyses to Rc grown seedlings, initially by expression analyses of *PIN1* in WT and *ceh1.* The analyses show similar *PIN1* transcript levels in *ceh1* and the WT seedlings (Fig. 6**A**). In contrast, the combined approaches of immunoblot and immunolocalization analyses confirmed a significant reduction in PIN1 protein levels in *ceh1* compared to WT seedlings (Fig. 6**B**-**C**). Specifically, immunolocalization clearly showed reduced PIN1 protein abundance in plasma membranes of xylem parenchyma cells (along tracheids), most notably in meristems of *ceh1* compared to WT seedling, albeit with an unchanged polarity (Fig. 6**C**). These data support the earlier finding establishing the role of MEcPP in modulating PIN1 prote in abundance both in Rc and in white light grown seedlings (Jiang et al., 2018).

**Fig. 6.**
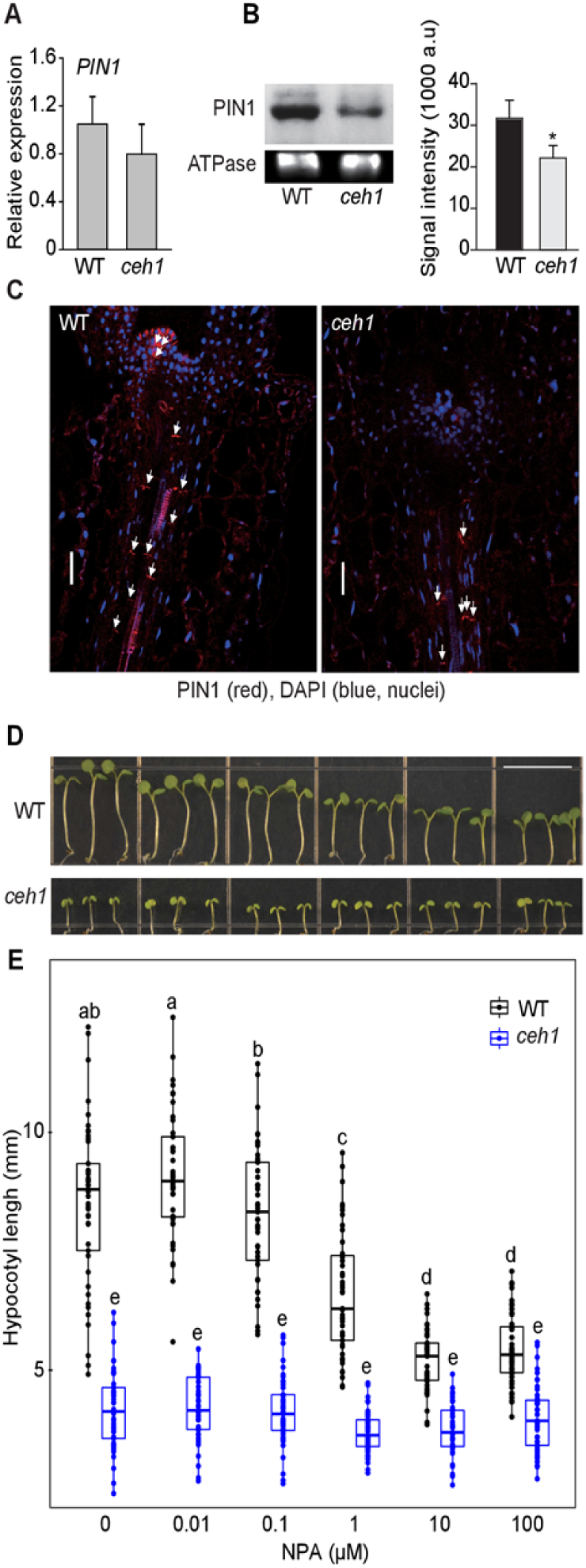
Altered auxin transport in ***ceh1***. (**A**) ***PIN1*** expression levels in 7-day-old WT and ***ceh1*** seedlings grown in Rc (15 μEm^−2^sec^−1^). Experiment was performed as described in Fig. 4A. Data are presented with three biological replicales and three technical replicates. (**B**) Immunobiots of PIN1 and ATPase as the protein loading control, and the signal quantification of PIN1 In 7-day-old WT and ***ceh1*** seedlings grown under Rc (15 μEm^−2^sec^−1^), Asterisk denotes significant difference as determinad by a two-tailed Student’s ***t*** tests. (**C**) Immunolocalization of PIN1 in 7-day-old WT and ***ceh1*** seedlings grown under Rc (15 μEm^−2^sec^−1^). Scale bar: 20 μm. (**D**) Representative Images of 7-day-old WT and ***ceh1*** seedlings grown under Rc (15 μEm^−2^sec^−1^) in the absence (0) and presenceof NPA. Scale bar: 1cm. (**E**) Quantification of hypocotyl length of seedlings from panel (D). Data are presented with 45 seedlings. Statistical analyses were carried out using Tukey’s HSD method, different letters indicate significant difference (*P* < 0.05),

The reduced levels of the major auxin transporter, PIN1, led us to examine the impact of varying concentrations of a general auxin polar transport inhibitor, 1-naphthylphthalamic acid (NPA) (Scanlon, 2003) on the hypocotyl growth of WT and *ceh1* seedlings grown in Rc (Fig. 6**D**-**E**). As expected, NPA application reduces WT hypocotyl growth in a dose-dependent manner. This contrasts with the lack of detectable response in *ceh1*, thereby confirming compromised auxin transport in the mutant.

### Ethylene regulates hypocotyl growth in ceh1

Comparative transcriptomic profiling of WT and *ceh1* seedlings grown in the Rc (15 μE m^−2^sec^−1^) revealed reduced levels of ethylene biosynthesis genes, *ACSs* (Table S1) in the mutant. This observation, in conjunction with the established crosstalk between ethylene and auxin (Yu et al., 2013; Sun et al., 2015; Das et al., 2016), prompted us to further investigate the potential function of ethylene in regulating *ceh1* hypocotyl growth. Initially, we performed qPCR analyses on ethylene biosynthesis genes to validate the original transcriptomic profile data (Table S1, and Fig. S4). The data shows that compared to WT seedlings there is a prominent reduction in the transcript levels of *ACS4* in the dark (≥2-fold) and the Rc (∼60-fold) grown *ceh1* seedlings, as well as a notable (3-to 10-fold depending on the gene) reduced expression of *ACS5, 6, 8* albeit solely in Rc-grown *ceh1* (Fig. 7**A**).

**Fig. 7.**
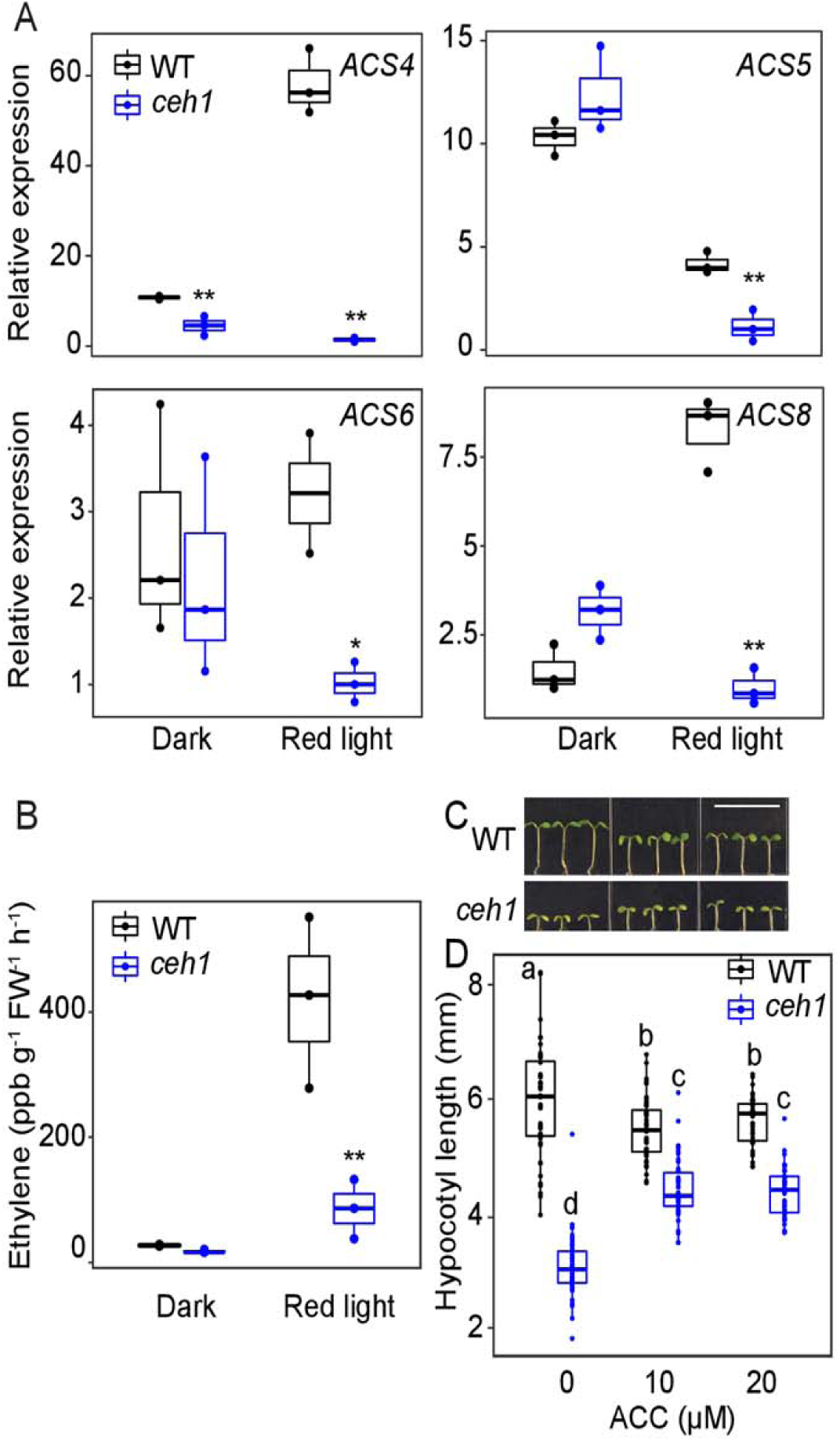
Ethylene regulates hypocotyl growth in *ceh1*. (**A**) Expression levels of *ACS4, 5, 6* and *8* in 7-day-old WT and *ceh1* seedlings grown in the dark and Rc (15 μEm^−2^sec^−1^). Experiment was performed as described in Fig. 4A. Data are presented with three biological replicates and three technical replicates. Statistical analyses were determined by a two-tailed Student’s *t* tests with a significance of *P* < 0.05 *, *P* < 0.01 **. (**B**) Ethylene levels in samples used in panel (A). (**C**) Representative images of 7-day-old WT and *ceh1* seedlings grown in the absence (0) and presence of ACC in the Rc (15 μEm^−2^sec^−1^). Scale bar: 1cm. (**D**) Quantification of hypocotyl length of seedlings from panel **(C).** Data are presented with 45 seedlings. Statistical analyses were carried out using Tukey’s HSD method, different letters indicate significant difference (*P* < 0,05).

Measurements of ethylene in these seedlings confirmed reduced levels (∼80%) of the hormone in Rc-grown *ceh1* compared to WT seedlings (Fig. 7**B**). This led us to examine hypocotyl growth of seedlings grown in the presence of varying concentrations of ethylene precursor, ACC (Fig. 7**C-D**). The data show suppression of WT hypocotyl growth at all concentrations examined, as opposed to equally enhanced hypocotyl growth in *ceh1* at both ACC concentrations (10 and 20 μM), an indication of saturation of growth response. Altogether, the data support MEcPP-mediated coordination of red-light signaling cascades with ethylene levels and ethylene regulation of hypocotyl growth.

### Hierarchy of ethylene and auxin signaling pathways

The partial recovery of *ceh1* hypocotyl growth by external application of auxin and ethylene, albeit to varying degrees, prompted us to genetically explore their potential interdependency and hierarchy of their respective growth regulatory actions in Rc grown seedlings. To address this, we applied ACC and IAA independently to single and double mutant lines of *ceh1* introgressed into auxin receptor mutant *tir1-1* (*ceh1/tir1-1*), and into single and double ethylene signaling mutants *ein3* and *eil1* (*ceh1/ein3, ceh1/eil1* and *ceh1/ein3 eil1*).

Analyses of hypocotyl lengths of Rc-grown WT, *ceh1, ceh1/tir1-1* and *tir1-1* seedlings in the absence and presence of ACC demonstrate TIR1-dependent growth promoting action of ACC in *tir1-1* and *ceh1/tir1-1* (Fig. 8**A**-**B**). We furthered these studies by applying ACC alone or together with NPA (Fig. 8**C-D**). Consistent with the earlier data, ACC treatment promoted *ceh1* hypocotyl growth, but less effectively when combined with auxin polar transport inhibitor, NPA (Fig. 8**C-D**).

**Fig. 8.**
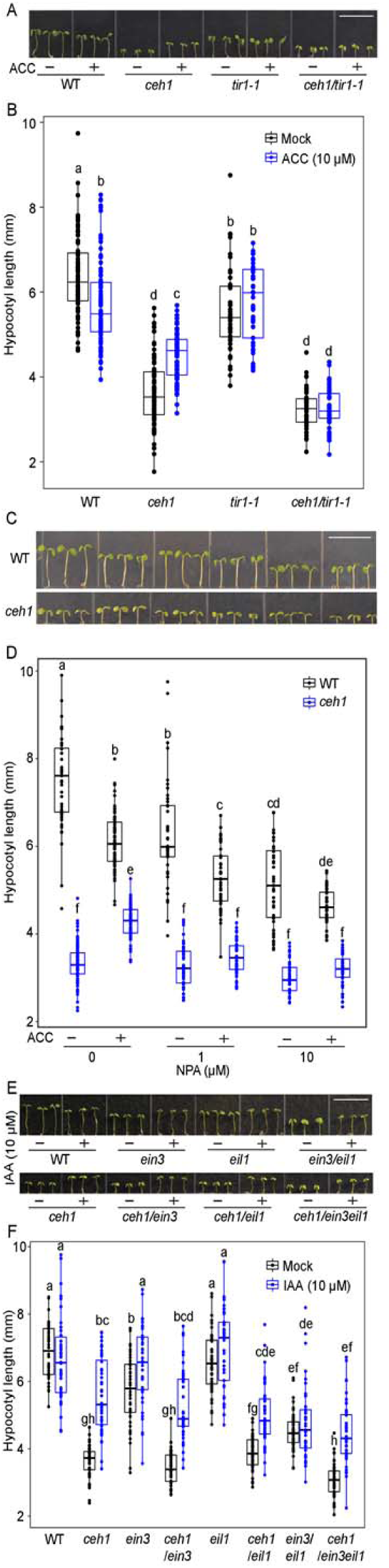
Auxin is epistatic to ethylene. (**A**) Representative images of 7-day-old WT. *ceh1, ceh1/tir1-1* and *tir1-1* seedlings grown in the Rc (15 μEm^−2^sec^−1^) in the absence (-) and presence (+) of ACC. (**C**) Representative images of 7-day-old WT and *ceh1* seedlings grown in the Rc (15 μEm^−2^sec^−1^) in the absence (-) and presence (+) of ACC/NPA alone or in combination. (**E**) Representative images of 7-day-old WT, *ceh1. ein3, ceh1/ein3, eil1. ceh1/eil1, ein3/eil1. ceh1/ein3eil1* seedlings grown in the Rc (15 μEm^−2^sec^−1^) in the absence (-) and presence (+) of IAA. (**B**) & (**D**) & (**F**) Quantification of hypocotyl length of seedlings from panel (**A**) & (**C**) & (**E**), respectively. Data are presented with 45 seedlings. Statistical analyses were carried out using Tukey’s HSD method, different letters indicate significant difference (*P* < 0.05). Scale bars: 1cm.

In parallel, we examined hypocotyl growth of Rc-grown WT, *ceh1, ein3, ceh1/ein3, eil1, ceh1/eil1, ein3eil1* and *ceh1*/*ein3eil1* seedlings in the presence and absence of externally applied IAA (Fig. 8**E-F**). Enhanced growth of *ceh1* hypocotyl in the presence of IAA irrespective of mutant backgrounds (single or double *ein3-eil1*) reaffirms growth-promoting function of auxin even in lines perturbed in ethylene signaling pathway.

This finding established the dependency of ethylene function on auxin signaling, and delineates the hierarchy of responses and positions auxin as being epistatic to ethylene signaling pathway.

## Discussion

An inherent feature of plant growth and development is the capacity to coordinate and integrate external cues with endogenous regulatory pathways through tightly regulated signaling cascades. Recent studies have identified retrograde signaling as a quintessential mode of cellular communication required for optimal organismal response to prevailing conditions. Here, we provide a coherent picture of how the stress-specific plastidial retrograde signaling metabolite (MEcPP) coordinates light and hormonal signaling circuitries to adjust growth to the most prevalent environmental cue, light conditions.

The simplified schematic model (Fig. 9) depicts MEcPP as the upstream signal coordinating and modulating drivers of growth, specifically through enhancing phyB protein abundance and the consequential reduction of auxin levels and distribution in conjunction with diminished ethylene content.

**Fig. 9.**
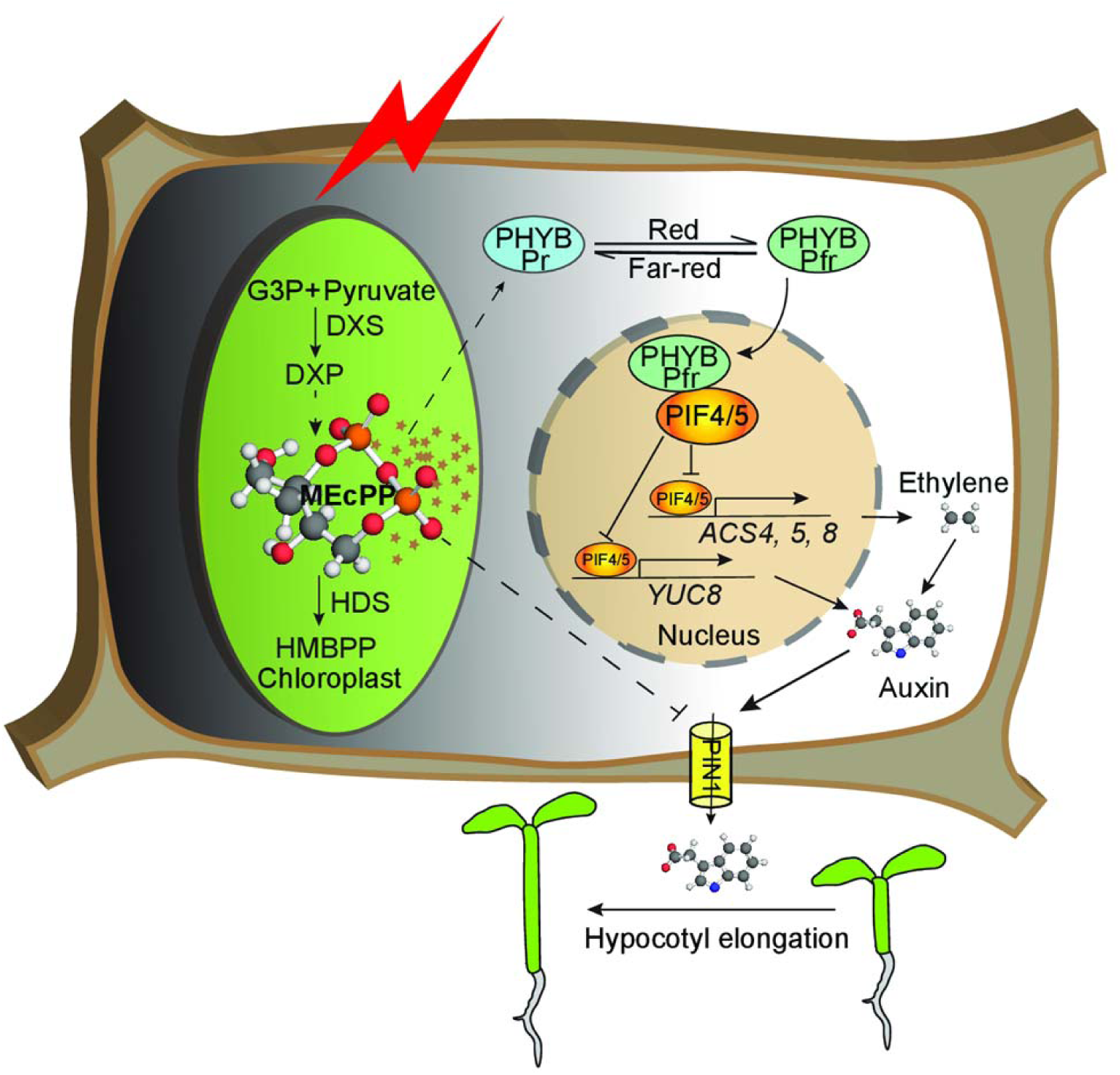
Schematic model depicting MEcPP as the integrator of growth regulating pathways Stress induction of MEcPP accumulation reduces expression of *PIF 4* and *5* and enhances abundance of phyB protein and the consequential orchestration of ethylene-auxin hierarchy to regulate growth.

The degradation of phyB is established to be through intermolecular transaction of this photoreceptor with PIF transcription factors (Ni et al., 2013), thereby supporting the prospect of significantly reduced *PIF4* and *5* transcript levels as the likely cause of enhanced phyB protein abundance in *ceh1* seedlings grown in Rc. Furthermore, reversion of *ceh1* stunted hypocotyls in *ceh1/phyB-9* confirms the key role of enhanced phyB protein abundance in growth retardation of the mutant, confirming the earlier finding using white light grown seedlings (Jiang et al., 2019)

The role of phyB in regulating growth is reported to be through repressing auxin response genes (Devlin et al., 2003; Halliday et al., 2009). The red light mediated reduction of auxin biosynthesis and signaling together with decreased hormone levels in *ceh1* supports phyB function in auxin regulation. In addition, reduced levels of PIN1 protein abundance as evidenced by immunoblot and immunolocalization assays suggest the regulatory role of phyB in controlling auxin transport via modulation of PIN1 protein levels. This notion is supported by the ineffectiveness of auxin transport inhibitor in modulating hypocotyl growth of Rc-grown *ceh1* seedlings.

Similar to auxin, reduction of ethylene levels, partly due to decreased transcript levels of the respective biosynthesis genes in Rc-grown *ceh1*, strongly supports the regulatory role of MEcPP-mediated induction of phyB in the process. Partial and differential recovery of *ceh1* hypocotyl growth under Rc in the presence of external auxin or ACC, identifies auxin as the key growth-regulating hormone under these experimental conditions. Moreover, measurement of hypocotyl growth of *ceh1* seedlings introgressed into auxin and ethylene signaling receptor mutants, places auxin epistatic to ethylene, and supports a one-directional control mechanism of ethylene-auxin interaction under Rc condition.

## Conclusions

Here, we reveal MEcPP-mediated enhanced abundance of PhyB, in part via suppression of *PIF4* and *5* expression, and the resulted reduced hypocotyl growth. We further establish MEcPP-mediated coordination of phytochrome B with auxin and ethylene signaling pathways, and the function of the collective signaling circuitries in regulation of hypocotyl growth of red-light grown seedlings. In addition, hormonal applications and pharmacological treatments support hierarchical functions of auxin and ethylene in regulating growth, with auxin being epistatic to ethylene.

In summary, this finding illustrates MEcPP-mediated coordination of light and hormonal signaling cascades to ultimately reprogram plant growth in responses to the light environment and further provides information on the functional hierarchy of these growth regulatory inputs. As such, this finding identifies plastids as the control hub of growth plasticity in response to environmental cues.

## METHODS

### Plant materials

The wild type seeding used here are the earlier reported Col-0 ecotype transformed with *HPL:LUC* constructs and used as the parent (P) for isolation of *ceh1* mutant (Xiao et al., 2012).

All experiments were performed with 7-day-old seedlings grown in15 μE m^−2^sec^−1^ continuous monochromatic light at 22 °C, unless specified otherwise. The *ein3eil1* double mutant is provided by Hongwei Guo (Southern University of Science and Technology); *DR5*-GFP is a gift from Mark Estelle (University of California, San Diego); *tir1-1* (CS3798) was ordered from ABRC.

### Light treatment

Surface-sterilized seeds were planted on half strength Murashige and Skoog medium (1/2 MS: 2.2 g/L Murashige and Skoog salts, 1g/L MES (2-(N-morpholino) ethanesulfonic acid, pH 5.7, and 8 g/L agar), stratified at 4 °C for 5 days, grown in 15 μE m^−2^sec^−1^ of monochromatic red, far-red and blue LEDs (Quantum Devices Snap-Lite) in a custom chamber at ∼ 22 °C for 7 days prior to hypocotyl measurement. Dark control experiments were performed by exposing seedlings to white light for 3 hours after stratification, and then wrapped the plates with 3 layers of aluminum foil, grown for 7 days before quantification of hypocotyl length. Each treatment was performed on three biological replicates, each replicate with 15 seedlings.

### Hypocotyl length measurement

7-day-old seedlings were scanned with an Epson flatbed scanner, hypocotyl length was measured using Image J.

### RNA isolation and RNA-Seq library construction

Total RNA was isolated from 7-day old seedlings grown in the dark and in Rc, using TRIzol (Life Technologies), the RNA quality and quantity were assessed by Nanodrop ND 1000 (Nanodrop technologies), 4 µg of qualified total RNA was used for RNA-Seq library preparation using Illumina’s TruSeq v1 RNA sample Preparation kit (RS-930-2002) with a low-throughput protocol following manufacturer’s instructions with modifications as described (Devisetty et al., 2014). Illumina’s 12 indices were used during adaptor ligation and library construction. The constructed libraries were size-selected using 1:1 volume of AMPure XP beads (Beckman Coulter, Brea CA). Size and quality of libraries were examined using Bioanalyzer 2100 (Agilent, Santa Clara, CA). The 12 libraries were quantified using Quant-iT™ PicoGreen® ds DNA Assay Kit (Invitrogen) and equally pooled in 1 lane of single end 50 bp sequencing in HiSeq 2000 machine (Illumina, San Diego, CA) at the QB3 facility at UC Berkeley.

### Quality filtering and alignment of RNA-Seq data

To ensure good read quality for downstream analysis, raw reads were pre-processed using FastX-tool kit software (http://hannonlab.cshl.edu/fastx_toolkit/) and custom Perl scripts. First, the de-multiplexed raw reads were filtered with fastq_quality_filter, parameters (−q 20, minimum quality score to keep: 20; −p 95, minimum percent of bases that must satisfy the quality score cut-off: 95). Next, reads with custom adapters were removed using a custom script. Quality of reads was examined before and after quality control with FastQC quality assessment software (http://www.bioinformatics.babraham.ac.uk/projects/fastqc/). Then Reads (1×50 bp) were mapped against the *Arabidopsis* representative_gene_model (TAIR10) using BWA v0.6.1-r104 (Li and Durbin, 2009) with parameters (-l 20) and SAMtools (Li et al., 2009). The resulting BAM files were used to calculate the read counts using a custom R script, and then the counts were used for differential gene expression analysis.

### Differential expression analysis of RNA-Seq data

The EdgeR Bioconductor package implemented in R was used to generate the pseudo-normalized counts for visualization and to carry out differential gene expression analysis (Robinson et al., 2010). Genes were kept for further analysis if read counts were greater than 1 count per million (cpm) in at least 3 of the 12 libraries. The EdgeR Generalized linear models (GLM) framework with explanatory variables of genotype and treatment allowed us to specify a design matrix estimating the effect of run number (batch) as a nuisance parameter. After fitting the model for our experiment, we defined contrasts between parent lines (WT) and mutant (*ceh1*) in red light and tested for significant expression differences using a likelihood ratio test (‘glmLRT’). *P*-values for the remaining genes were adjusted using Benjamini-Hochberg method for false discovery correction. Genes with an FDR-adjusted *P*-value less than or equal to 0.01 were identified as differentially expressed.

### Multi-dimensional scaling (MDS) plot

A multi-dimensional scaling (MDS) plot was generated in edgeR to analyze relationship between samples. Distance between each pair of RNA-seq profiles corresponded to the average (root-mean-square) of absolute logFC between each pair of samples.

### GO Term enrichment

Goseq package in R (Young et al., 2010) was used to identify enriched Gene Ontology (GO) terms (mainly biochemical process) in the differentially expressed gene list.

### Hormones and chemical treatments

Surface-sterilized seeds were planted on 1/2 MS, stratified at 4 °C for 3 days, germinated under continuous red light at 15 μmol m^−2^sec^−1^ for 2 days and subsequently transformed to 1/2 MS medium with 1 g/L MES (2-(N-morpholino) ethanesulfonic acid) in combination with hormones or chemicals. These plates were vertically placed in continuous red light for 5 extra days before hypocotyl measurements. IAA, ACC, auxinole and NPA were dissolved in ethanol, water, DMSO and DMSO, respectively. The corresponding solvents were used as control treatment (mock) for the respective experiments.

### MEcPP and hormone measurements

Quantification of SA, JA,ABA and IAA was carried out by gas chromatography-mass spectrometry (GC-MS), using dihydro-JA and deuterated SA and ABA, and IAA as internal standard, as previously described (Savchenko et al., 2010). MEcPP extraction and quantification was performed as previously described (Jiang et al., 2019).

### Microscopy

Confocal fluorescence imaging was performed using Zeiss LSM 710. GFP signal was examined in 7-day-old *DR5*-GFP and *ceh1/DR5*-GFP seedlings grown on 1/2 MS in Rc (15 μE m^−2^sec^−1^).

### Immunolocalization of PIN1

Immunolocalization of PIN1 was performed using anti-PIN1 monoclonal primary antibody and FITC anti mouse secondary antibody as previously described (Jiang et al., 2018).

### Quantitative RT-PCR

Total RNAs were isolated from 7-day-old seedlings grown in the Rc using TRIzol (Life Technologies), treated with DNase to eliminate DNA contamination. 1 μg total RNA was reverse transcribed into cDNA using SuperScript III (Invitrogen). *At4g26410* was used to normalize target gene expressions. Gene-specific primers were designed using QuantPrime q-PCR primer design tool (http://www.quantprime.de/) and are listed (Supplemental Table 2). Each experiment was performed with three biological replicates and three technical replicates.

### Protein extraction and immunoblot analyses

For protein extraction 7-day-old seedlings were collected, ground with liquid nitrogen, homogenized in extraction buffer (10 mM Hepes, pH 7.6, 1 M Sucrose, 5 mM KCl, 5 mM MgCl2, 5 mM EDTA, 14 mM 2-ME, 0.4% Triton X-100, 0.4 mM PMSF, 20 μM MG132, 20 μM MG115, and Proteinase Inhibitor), centrifuged at 10,000 rpm for 10 min at 4 °C, supernatants were transferred to new tubes as total proteins. Then the proteins were separated on 7.5 % SDS-PAGE gel, transferred to PVDF membranes. Blots were probed with B1+B7 (1:500) primary antibodies obtained from Peter Quail lab. The secondary was anti-mouse horseradish peroxidase (HRP) (KPL, catalog no. 074-1806) (1:10000). Immunoblots for PIN1 protein were performed as previously described (Jiang et al., 2018) using anti PIN1 monoclonal antibody (1:100) primary antibody and secondary anti-mouse Horseradish peroxidase (HRP) (1:3000). Chemiluminescent reactions were performed using the Pierce ECL Western Blotting Substrate following the manufacturer’s instructions. The excessive substrate was removed from membranes before placing them between two plastic sheets to develop with X-ray, and subsequently scanned with Epson Perfection V600 Photo Scanner.

### Statistical analyses

All experiments were performed with at least three biological replicates. Data are mean ± standard deviation (SD). The Statistical test was performed using library agricolae, Tukey’s HSD test method in R with a significance of *P* < 0.05 (Bunn, 2008). We have specified the method we used for statistical test in all figure legends.

## Acknowledgements

We would like to thank Mr. Derrick R. Hicks (University of Washington) for providing images of ethylene, IAA and MEcPP stick chemical structures depicted in our model. We also would like to thank Prof. Meng Chen and Dr. Yongjian Qiu for providing the red light chamber and reagents for our experiments, and Dr. Peter Quail for generously providing us with the phyB antibody. We would like to thank Jacob North for all his efforts towards seedling preparation. We are thankful to Dr. Geoffrey Benn for performing the statistical analyses using R program.

## Author Contributions

J.J. and K.D. designed the study, J.J., Y.X., H.C. W.H., U.D., H.K., and F.D. performed the experiments, L.Z performed the statistical analyses, J.M. and K.P. provided experimental tools and K.D. wrote the manuscript.

## Parsed Citations

Bjornson, M., Balcke, G.U., Xiao, Y., de Souza, A., Wang, J.Z., Zhabinskaya, D., Tagkopoulos, I., Tissier, A., and Dehesh, K. (2017). Integrated omics analyses of retrograde signaling mutant delineate interrelated stress-response strata. Plant J 91, 70–84.

Bunn, A.G. (2008). Adendrochronology program library in R (dplR). Dendrochronologia 26, 115–124.

Chai, T., Zhou, J., Liu, J., and Xing, D. (2015). LSD1 and HY5 antagonistically regulate red light induced-programmed cell death in Arabidopsis. Front Plant Sci 6, 292.

Chenge-Espinosa, M., Cordoba, E., Romero-Guido, C., Toledo-Ortiz, G., and Leon, P. (2018). Shedding light on the methylerythritol phosphate (MEP)-pathway: long hypocotyl 5 (HY5)/phytochrome-interacting factors (PIFs) transcription factors modulating key limiting steps. Plant J 96, 828–841.

Das, D., St Onge, K.R., Voesenek, L.A., Pierik, R., and Sasidharan, R. (2016). Ethylene- and shade-induced hypocotyl elongation share transcriptome patterns and functional regulators. Plant Physiol.

De Grauwe, L., Vandenbussche, F., Tietz, O., Palme, K., and Van Der Straeten, D. (2005). Auxin, ethylene and brassinosteroids: tripartite control of growth in the Arabidopsis hypocotyl. Plant & cell physiology 46, 827–836.

de Wit, M., Lorrain, S., and Fankhauser, C. (2014). Auxin-mediated plant architectural changes in response to shade and high temperature. Physiologia plantarum 151, 13–24.

Devisetty, U.K., Covington, M.F., Tat, A.V., Lekkala, S., and Maloof, J.N. (2014). Polymorphism identification and improved genome annotation of Brassica rapa through Deep RNAsequencing. G3 (Bethesda) 4, 2065–2078.

Devlin, P.F., Yanovsky, M.J., and Kay, S.A. (2003). Agenomic analysis of the shade avoidance response in Arabidopsis. Plant physiology 133, 1617–1629.

Franklin, K.A., Lee, S.H., Patel, D., Kumar, S.V., Spartz, A.K., Gu, C., Ye, S., Yu, P., Breen, G., Cohen, J.D., Wigge, P.A., and Gray, W.M. (2011). Phytochrome-interacting factor 4 (PIF4) regulates auxin biosynthesis at high temperature. Proceedings of the National Academy of Sciences of the United States of America 108, 20231–20235.

Galweiler, L., Guan, C., Muller, A., Wisman, E., Mendgen, K., Yephremov, A., and Palme, K. (1998). Regulation of polar auxin transport by AtPIN1 in Arabidopsis vascular tissue. Science 282, 2226–2230.

Geldner, N., Friml, J., Stierhof, Y.D., Jurgens, G., and Palme, K. (2001). Auxin transport inhibitors block PIN1 cycling and vesicle trafficking. Nature 413, 425–428.

Gonzalez-Cabanelas, D., Wright, L.P., Paetz, C., Onkokesung, N., Gershenzon, J., Rodriguez-Concepcion, M., and Phillips, M.A. (2015). The diversion of 2-C-methyl-D-erythritol-2,4-cyclodiphosphate from the 2-C-methyl-D-erythritol 4-phosphate pathway to hemiterpene glycosides mediates stress responses in Arabidopsis thaliana. The Plant journal : for cell and molecular biology 82, 122–137.

Halliday, K.J., Martinez-Garcia, J.F., and Josse, E.M. (2009). Integration of light and auxin signaling. Cold Spring Harbor perspectives in biology 1, a001586.

Hayashi, K., Tan, X., Zheng, N., Hatate, T., Kimura, Y., Kepinski, S., and Nozaki, H. (2008). Small-molecule agonists and antagonists of F-box protein-substrate interactions in auxin perception and signaling. Proceedings of the National Academy of Sciences of the United States of America 105, 5632–5637.

Hayashi, K., Neve, J., Hirose, M., Kuboki, A., Shimada, Y., Kepinski, S., and Nozaki, H. (2012). Rational design of an auxin antagonist of the SCF(TIR1) auxin receptor complex. ACS Chem Biol 7, 590–598.

Hornitschek, P., Kohnen, M.V., Lorrain, S., Rougemont, J., Ljung, K., Lopez-Vidriero, I., Franco-Zorrilla, J.M., Solano, R., Trevisan, M., Pradervand, S., Xenarios, I., and Fankhauser, C. (2012a). Phytochrome interacting factors 4 and 5 control seedling growth in changing light conditions by directly controlling auxin signaling. Plant J 71, 699–711.

Hornitschek, P., Kohnen, M.V., Lorrain, S., Rougemont, J., Ljung, K., Lopez-Vidriero, I., Franco-Zorrilla, J.M., Solano, R., Trevisan, M., Pradervand, S., Xenarios, I., and Fankhauser, C. (2012b). Phytochrome interacting factors 4 and 5 control seedling growth in changing light conditions by directly controlling auxin signaling. The Plant journal : for cell and molecular biology 71, 699–711.

J. Jensen, P., P. Hangarter, R., and Estelle, M. (1998). Auxin Transport Is Required for Hypocotyl Elongation in Light-Grown but Not Dark-Grown Arabidopsis. plant Physiol 116, 455–462.

Jiang, J., Zeng, L., Ke, H., De La Cruz, B., and Dehesh, K. (2019). Orthogonal regulation of phytochrome B abundance by stress-specific plastidial retrograde signaling metabolite. Nature communications 10, 2904.

Jiang, J., Rodriguez-Furlan, C., Wang, J.Z., de Souza, A., Ke, H., Pasternak, T., Lasok, H., Ditengou, F.A., Palme, K., and Dehesh, K. (2018). Interplay of the two ancient metabolites auxin and MEcPP regulates adaptive growth. Nature communications 9, 2262.

L.-C. Wang, K., Li, H., and R. Ecker, J. (2002). Ethylene Biosynthesis and Signaling Networks. The plant cell, 131–151.

Leivar, P., and Quail, P.H. (2011). PIFs: pivotal components in a cellular signaling hub. Trends in plant science 16, 19–28.

Leivar, P., and Monte, E. (2014). PIFs: Systems Integrators in Plant Development. Plant Cell 26, 56–78.

Leivar, P., Monte, E., Cohn, M.M., and Quail, P.H. (2012). Phytochrome signaling in green Arabidopsis seedlings: impact assessment of a mutually negative phyB-PIF feedback loop. Molecular plant 5, 734–749.

Lemos, M., Xiao, Y., Bjornson, M., Wang, J.Z., Hicks, D., Souza, A., Wang, C.Q., Yang, P., Ma, S., Dinesh-Kumar, S., and Dehesh, K. (2016). The plastidial retrograde signal methyl erythritol cyclopyrophosphate is a regulator of salicylic acid and jasmonic acid crosstalk. J Exp Bot 67, 1557–1566.

Li, H., and Durbin, R. (2009). Fast and accurate short read alignment with Burrows-Wheeler transform. Bioinformatics 25, 1754–1760.

Li, H., Handsaker, B., Wysoker, A., Fennell, T., Ruan, J., Homer, N., Marth, G., Abecasis, G., Durbin, R., and Genome Project Data

Processing, S. (2009). The Sequence Alignment/Map format and SAMtools. Bioinformatics 25, 2078–2079.

Liang, X., Wang, H., Mao, L., Hu, Y., Dong, T., Zhang, Y., Wang, X., and Bi, Y. (2012). Involvement of COP1 in ethylene- and light-regulated hypocotyl elongation. Planta 236, 1791–1802.

Morelli, G., and Ruberti, I. (2002). Light and shade in the photocontrol of Arabidopsis growth. Trends in plant science 7, 399–404.

Nagy, F., and Schafer, E. (2002). Phytochromes control photomorphogenesis by differentially regulated, interacting signaling pathways in higher plants. Annual review of plant biology 53, 329–355.

Negi, S., Sukumar, P., Liu, X., Cohen, J.D., and Muday, G.K. (2010). Genetic dissection of the role of ethylene in regulating auxin-dependent lateral and adventitious root formation in tomato. Plant J 61, 3–15.

Ni, W., Xu, S.L., Chalkley, R.J., Pham, T.N., Guan, S., Maltby, D.A., Burlingame, A.L., Wang, Z.Y., and Quail, P.H. (2013). Multisite light-induced phosphorylation of the transcription factor PIF3 is necessary for both its rapid degradation and concomitant negative feedback modulation of photoreceptor phyB levels in Arabidopsis. The Plant cell 25, 2679–2698.

Nozue, K., Harmer, S.L., and Maloof, J.N. (2011). Genomic Analysis of Circadian Clock-, Light-, and Growth-Correlated Genes Reveals PHYTOCHROME-INTERACTING FACTOR5 as a Modulator of Auxin Signaling in Arabidopsis. Plant Physiol 156, 357–372.

Nozue, K., Devisetty, U.K., Lekkala, S., Mueller-Moule, P., Bak, A., Casteel, C.L., and Maloof, J.N. (2018). Network Analysis Reveals a Role for Salicylic Acid Pathway Components in Shade Avoidance. Plant Physiol 178, 1720–1732.

Pedmale, U.V., Huang, S.C., Zander, M., Cole, B.J., Hetzel, J., Ljung, K., Reis, P.A.B., Sridevi, P., Nito, K., Nery, J.R., Ecker, J.R., and Chory, J. (2016). Cryptochromes Interact Directly with PIFs to Control Plant Growth in Limiting Blue Light. Cell 164, 233–245.

Quail, P.H. (2002). Photosensory perception and signalling in plant cells: new paradigms? Current opinion in cell biology 14, 180–188.

Rausenberger, J., Hussong, A., Kircher, S., Kirchenbauer, D., Timmer, J., Nagy, F., Schafer, E., and Fleck, C. (2010). An Integrative Model for Phytochrome B Mediated Photomorphogenesis: From Protein Dynamics to Physiology. Plos One 5.

Robinson, M.D., McCarthy, D.J., and Smyth, G.K. (2010). edgeR: a Bioconductor package for differential expression analysis of digital gene expression data. Bioinformatics 26, 139–140.

Ruzicka, K., Ljung, K., Vanneste, S., Podhorska, R., Beeckman, T., Friml, J., and Benkova, E. (2007). Ethylene regulates root growth through effects on auxin biosynthesis and transport-dependent auxin distribution. The Plant cell 19, 2197–2212.

Savchenko, T., Walley, J.W., Chehab, E.W., Xiao, Y., Kaspi, R., Pye, M.F., Mohamed, M.E., Lazarus, C.M., Bostock, R.M., and Dehesh, K. (2010). Arachidonic acid: an evolutionarily conserved signaling molecule modulates plant stress signaling networks. The Plant cell 22, 3193–3205.

Scanlon, M.J. (2003). The polar auxin transport inhibitor N-1-naphthylphthalamic acid disrupts leaf initiation, KNOX protein regulation, and formation of leaf margins in maize. Plant Physiol 133, 597–605.

Sellaro, R., Pacin, M., and Casal, J.J. (2012). Diurnal dependence of growth responses to shade in Arabidopsis: role of hormone, clock, and light signaling. Molecular plant 5, 619–628.

Smalle, J., Haegman, M., Kurepa, J., Van Montagu, M., and Van Der Straeten, D. (1997). Ethylene can stimulate Arabidopsis hypocotyl elongation in the light. P Natl Acad Sci USA 94, 2756–2761.

Stepanova, A.N., Yun, J., Likhacheva, A.V., and Alonso, J.M. (2007). Multilevel interactions between ethylene and auxin in Arabidopsis roots. Plant Cell 19, 2169–2185.

Sun, J., Ma, Q., and Mao, T. (2015). Ethylene Regulates the Arabidopsis Microtubule-Associated Protein WAVE-DAMPENED2-LIKE5 in Etiolated Hypocotyl Elongation. Plant Physiol 169, 325–337.

Swarup, R., Perry, P., Hagenbeek, D., Van Der Straeten, D., Beemster, G.T.S., Sandberg, G., Bhalerao, R., Ljung, K., and Bennett, M.J. (2007). Ethylene upregulates auxin biosynthesis in Arabidopsis seedlings to enhance inhibition of root cell elongation. Plant Cell 19, 2186–2196.

Tanaka, S., Nakamura, S., Mochizuki, N., and Nagatani, A. (2002a). Phytochrome in cotyledons regulates the expression of genes in the hypocotyl through auxin-dependent and -independent pathways. Plant & cell physiology 43, 1171–1181.

Tanaka, S.I., Nakamura, S., Mochizuki, N., and Nagatani, A. (2002b). Phytochrome in cotyledons regulates the expression of genes in the hypocotyl through auxin-dependent and -independent pathways. Plant and Cell Physiology 43, 1171–1181.

Tian, Q., Uhlir, N.J., and Reed, J.W. (2002). Arabidopsis SHY2/IAA3 inhibits auxin-regulated gene expression. The Plant cell 14, 301–319.

Vandenbussche, F., Vaseva, I., Vissenberg, K., and Van Der Straeten, D. (2012). Ethylene in vegetative development: a tale with a riddle. The New phytologist 194, 895–909.

Vandenbussche, F., Smalle, J., Le, J., Saibo, N.J., De Paepe, A., Chaerle, L., Tietz, O., Smets, R., Laarhoven, L.J., Harren, F.J., Van Onckelen, H., Palme, K., Verbelen, J.P., and Van Der Straeten, D. (2003). The Arabidopsis mutant alh1 illustrates a cross talk between ethylene and auxin. Plant Physiol 131, 1228–1238.

Walley, J., Xiao, Y., Wang, J.Z., Baidoo, E.E., Keasling, J.D., Shen, Z., Briggs, S.P., and Dehesh, K. (2015). Plastid-produced interorgannellar stress signal MEcPP potentiates induction of the unfolded protein response in endoplasmic reticulum. Proceedings of the National Academy of Sciences of the United States of America 112, 6212–6217.

Wang, J.Z., Li, B., Xiao, Y., Ni, Y., Ke, H., Yang, P., de Souza, A., Bjornson, M., He, X., Shen, Z., Balcke, G.U., Briggs, S.P., Tissier, A., Kliebenstein, D.J., and Dehesh, K. (2017a). Initiation of ER Body Formation and Indole Glucosinolate Metabolism by the Plastidial Retrograde Signaling Metabolite, MEcPP. Molecular plant 10, 1400–1416.

Wang, J.Z., Li, B.H., Xiao, Y.M., Ni, Y., Ke, H.Y., Yang, P.Y., de Souza, A., Bjornson, M., He, X., Shen, Z.X., Balcke, G.U., Briggs, S.P., Tissier, A., Kliebenstein, D.J., and Dehesh, K. (2017b). Initiation of ER Body Formation and Indole Glucosinolate Metabolism by the Plastidial Retrograde Signaling Metabolite, MEcPP. Molecular plant 10, 1400–1416.

Xiao, Y., Savchenko, T., Baidoo, E.E., Chehab, W.E., Hayden, D.M., Tolstikov, V., Corwin, J.A., Kliebenstein, D.J., Keasling, J.D., and Dehesh, K. (2012). Retrograde signaling by the plastidial metabolite MEcPP regulates expression of nuclear stress-response genes. Cell 149, 1525–1535.

Yang, S.F., and Hoffman, N.E. (1984). Ethylene Biosynthesis and Its Regulation in Higher-Plants. Annu Rev Plant Phys 35, 155–189.

Young, M.D., Wakefield, M.J., Smyth, G.K., and Oshlack, A. (2010). Gene ontology analysis for RNA-seq: accounting for selection bias. Genome Biol 11, R14.

Yu, X., Liu, H., Klejnot, J., and Lin, C. (2010). The Cryptochrome Blue Light Receptors. The arabidopsis book 8, e0135.

Yu, Y., Wang, J., Zhang, Z., Quan, R., Zhang, H., Deng, X.W., Ma, L., and Huang, R. (2013). Ethylene promotes hypocotyl growth and HY5 degradation by enhancing the movement of COP1 to the nucleus in the light. PLoS genetics 9, e1004025.

Zhao, Y. (2012). Auxin biosynthesis: a simple two-step pathway converts tryptophan to indole-3-acetic acid in plants. Mol Plant 5, 334–338.

Zhong, S., Shi, H., Xue, C., Wang, L., Xi, Y., Li, J., Quail, P.H., Deng, X.W., and Guo, H. (2012). A molecular framework of light-controlled phytohormone action in Arabidopsis. Curr Biol 22, 1530–1535.

